# Distinguishing and employing sister species of fish in assessment of stream quality

**DOI:** 10.1101/2021.08.12.456153

**Authors:** Fred Van Dyke, Benjamin W. van Ee, Seth Harju, Joshua W. Budi, Thomas B. Sokolowski, Brian Keas

**Author notes:** Current address: 6981 Rainbow Lake Road NE, Mancelona, Michigan, United States of America. Current address: 924 Kellogg Ave, Unit 2, Ames, Iowa, United States of America (Joshua W. Budi). Current address: 3400 Henley St., Glenview, Illinois, United States of America (Thomas B. Sokolowski). Corresponding author (FVD). **Author Contributions** FVD conceptualized, administered, supervised, and obtained funding for the study, and was lead and corresponding author; JB, TS were field and laboratory investigators; BVE conceptualized, administered, and supervised genetic analyses, interpretation and additional laboratory work subsequent to field data collection; SH analyzed non-genetic data; FVD, BVE, JB, TS, SH wrote and edited the manuscript; BK contributed to manuscript preparation and editing.

## Abstract

Biological indicators (bioindicators) can be individual species or species groups used to assess habitat quality. Unfortunately, conservationists often lack information on species distribution, how to differentiate between similar species, and environmental conditions associated with the presence of a species. We addressed these problems using two “sister” species of fish, the Mottled Sculpin (*Cottus bairdii*) and the Slimy Sculpin (*Cottus cognatus*), as stream quality indicators in the Manistee River watershed in the Huron-Manistee National Forests in Michigan, USA. We determined the abundance and distribution of these species and related their presence to concurrent in-stream measurements of temperature, dissolved oxygen, pH, conductivity, turbidity, and stream quality score based on macroinvertebrate diversity. To be certain of identification, we sequenced the Cytochrome c Oxidase Subunit I (CO1) molecular marker for specimens and used it as a DNA barcode to determine a specimen’s species. Cladistic analyses of CO1 unambiguously supported recognition of Mottled Sculpin and Slimy Sculpin as distinct species, confirming initial 87.5% correct identification using morphological characteristics, with uncertainty limited to juvenile fish. Field determinations increased to 100% correct identification as investigators gained more experience. Both species were most abundant in headwater regions, decreased downstream, and were sympatric at several locations. Mottled Sculpin were more likely to be found at stream locations with lower conductivity, pH, and stream quality scores, whereas Slimy Sculpin presence was more strongly associated higher levels of DO and lower levels of turbidity. Such findings are important because Mottled Sculpin are a designated management indicator species of the US Forest Service in the Huron-Manistee National Forests, but may be ineffective as a habitat quality indicator when used alone. Concurrent use of Mottled Sculpin and Slimy Sculpin as a management indicator sister-species complex could allow sufficient landscape coverage to permit habitat assessment if species-specific differences in environmental tolerances are precisely determined.

## Introduction

Biotic indices are widely used for monitoring the health of ecosystems and the quality of habitat [1]. Multi-metric and multi-species indices offer comprehensive assessments of environmental conditions and have been used with success, especially in aquatic environments. For these aquatic environmental evaluations, fish are an important species group. Indeed, one of the oldest and most widely used aquatic assessment tools, the Index of Biological Integrity (IBI), is entirely fish-based, with metrics including species composition, richness, tolerance, hybridization, trophic measures, health condition, age structure, growth and recruitment [2, 3]. More recently, multi-metric assessment indices, including multi-metric vegetation, invertebrate, and fish-based assessments, have been widely employed [4, 5]. Multi-metric indicators have also become increasingly quantified, and the growing use of Indicator Species Analysis (ISA) permits the quantification of the relative value of species or species groups as indicators [6, 7]. ISA is an appealing approach because it can produce an independently calculated value for each species (IndVal), be used with any species data set that accounts for the abundance and frequency of each species, and be applied to any sampling typology. More advanced use of ISA and IndVal has been made with applications designed to achieve Threshold Indicator Analysis (TITAN analysis) that calculates species and community thresholds with relation to different environmental variables or pollutants. However, TITAN has not been able to detect thresholds associated with abrupt changes in rates (slopes) or direction; rather, it has been more successful and predictive with Step Function responses than other forms of ecological responses [8].

Despite the potential of increased comprehensiveness from multiple species, which include the promise of greater and more precise quantification, a single species or single group of species whose function, population, or presence can be used to determine ecosystem performance or environmental change can also act as an effective biological indicator, or “indicator species” [3, 9]. Such indicators can provide, especially in single species, easy and cost-effective tools for short- and long-term monitoring of environmental and ecosystem integrity [10]—an important consideration when funding is limited and when scientists are asked to give reliable and rapid judgments about the quality of aquatic habitat at the request of government agencies or private institutions [1]. To be effective, indicator species should exhibit a narrow ecological range, rapid response to environmental change, well-defined taxonomy for reliable identification, wide distribution, and low-cost sampling [11]. Indicator species that meet these criteria can provide value to conservationists as surrogates representing the presence or abundance of other species or of environmental conditions. Properly employed, they can offer rapid and relatively simple techniques for habitat and species inventory in complex landscapes [12]. However, despite the ease and simplicity of employing a single species, confusion can still occur regarding the use and interpretation of indicator species presence or abundance. There are at least seven kinds of indicator species described in the scientific literature, each “indicating” something different [13, 14]. Indicator species may be used as “indicators” of the presence of a particular habitat or ecosystem (“composition indicators”) or of the condition of a habitat, community, or ecosystem (“condition indicators”). In some cases the same species or species group is used for both [15].

One type of indicator species used to establish species-site associations is the “management indicator species” (MIS), defined as “an organism whose characteristics, such as presence or absence, population density, dispersion, or reproductive success are used as an index of attributes [usually of the surrounding environment or habitat] too difficult, inconvenient, or expensive to measure” [16]. Thorough knowledge of environmental tolerances and preferences of these species is essential if MIS are to have value in habitat assessment and facilitate judicious and rapid decisions at minimal cost [13]—all elements which are critical for effective biodiversity conservation. The use of indicator species to characterize qualitative environmental preferences can then be associated with specific sites or environmental conditions. Such use has led to their designation as “diagnostic” species [17] or “sentinel species,” which provide early warning of environmental danger or hazard [18]. In both US and Canadian environmental agencies, individual species have been used as indicator or sentinel species.

An alternative to the use of individual species or metrics developed from unrelated species is the employment of closely related or “sister species.” Species relatedness can be an advantage in environmental assessment because both inter-specific competition and small - but important - differences in habitat quality could affect occurrence and abundance, and multiple mechanisms may interact simultaneously [19]. One pair of sister species that might offer insight into stream habitat assessment is the Mottled Sculpin (*Cottus bairdii*, Girard 1850) and Slimy Sculpin (*Cottus cognatus*, Richardson 1836). Both are potentially attractive as MIS and sentinel species because of their stable, secure, and localized (but widespread) populations ([20, 21], coupled with their occurrence in cold-water systems [18, 22]. Site sensitivity is enhanced because both can speciate in different but proximate streams ([23, 24, 25] and, like other species of sculpins, lack a swim bladder and inhabit benthic habitats, reducing their mobility and contributing to strong site fidelity, with high susceptibility to local change [18, 26, 27, 28].

Collectively, these qualities make Mottled Sculpin and Slimy Sculpin relatively site-specific MIS and sentinel species, and, in theory, diagnostic of high-quality stream conditions, such as high oxygenation levels, low organic content, and neutral to slightly alkaline pH. Sculpin species have an estimated upper lethal water limit between 23 and 25 °C [29, 30], but Slimy Sculpin are rarely found in waters with sustained temperatures >19 °C, and Mottled Sculpin in some western North American streams have shown even colder upper thermal limits [25]. Other abiotic variables, including levels of dissolved oxygen (DO), pH, and turbidity may influence sculpin presence, abundance, and ecology [31, 32, 33]. Both species are sensitive to toxic metals [34, 35], and Slimy Sculpin have shown response to environmental contaminants associated with agriculture [36, 37], coal mining [38], pulp and paper operations [39], and sewage [40].

The Slimy Sculpin has been chosen as a sentinel species in Canada because of its sensitivity to environmental pollutants, and in the United States, the Mottled Sculpin was declared an MIS of stream quality by the US Forest Service (USFS) for the Huron-Manistee National Forests (HMNF) in Michigan in 2013 [41]. Both species occur in Michigan’s Manistee River, where the HMNF has significant responsibilities for much of the area as the principal land use manager in the approximately 4,660 km^2^ watershed. Some sections of the Manistee River have been variously designated as a Blue Ribbon Trout Stream, a National Recreational River, and a National Wild and Scenic River. Portions of the river also flow through lands of the federally recognized Little River Band of Ottawa Indians (LRBOI), making the Manistee a river of high cultural significance. Given the importance of this river, combined with designation of the Mottled Sculpin as an MIS and the concurrent presence of the environmentally sensitive Slimy Sculpin, we sought to determine whether these species would be reliable indicators of environmental conditions in the local landscape context of the Huron-Manistee National Forests as well as non-HMNF lands in the upper Manistee River watershed.

Criteria established by the USFS for the selection of the Mottled Sculpin as one of six MIS for the HMNF included known distribution, well-documented response to stream alteration, and an important ecological role in the habitat [41]— characteristics that can also apply to the Slimy Sculpin [18]. The similarity of environmental criteria for these two species is particularly relevant when used as indicators, as other species of sculpin may have different water quality tolerances [31]. Environmental correspondence is accompanied by morphological similarity, which could limit the value of either species acting alone as a biological indicator if they could be misidentified. Both species are comparable in size, have large heads, stout bodies, wide gapes, and are dorsoventrally compressed [42] (Fig 1). Various morphological traits have been proposed to separate the two species [43, 44], but such checks and measurements can lead to uncertain discrimination in the field, and genetic analysis may be needed to resolve species identification [45,18] and to determine the relationships between morphological and molecular differences, which have not been thoroughly investigated.

**Fig 1.**
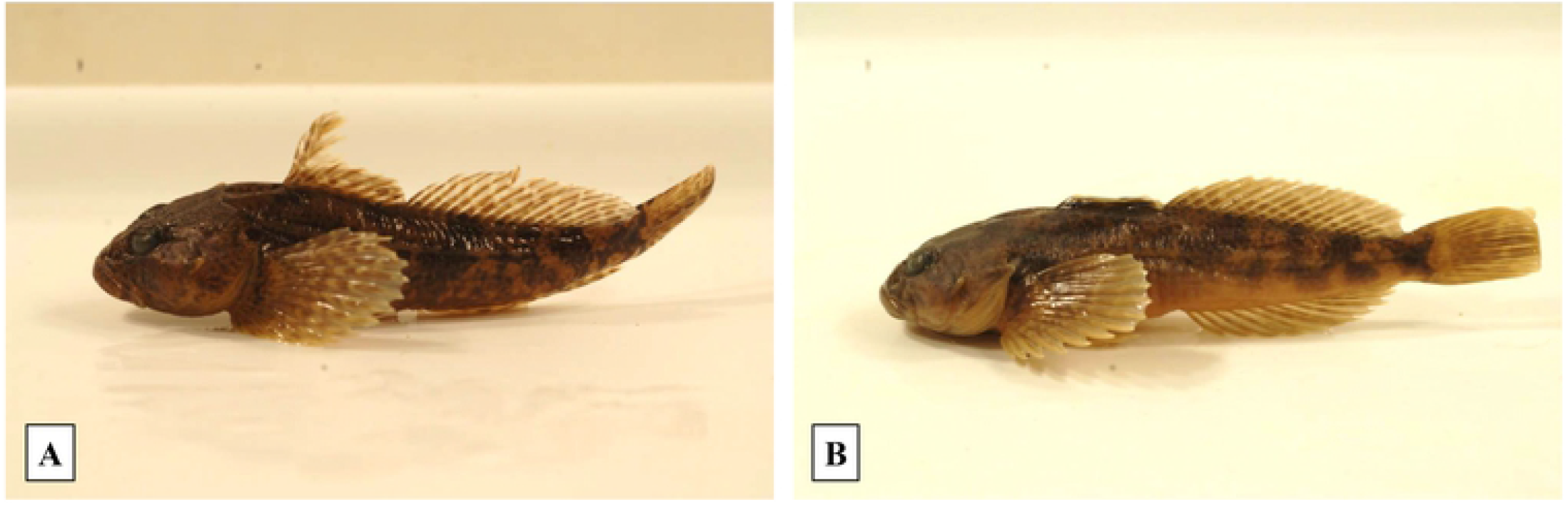
Mottled Sculpin (Cottus bairdii) (A) and Slimy Sculpin (Cottus cognatus) (B) are morphologically similar. One morphological difference is the number of pelvic fin rays – normally four in the Mottled Sculpin and three in the Slimy Sculpin. Sculpin photos courtesy of U. S. Fish and Wildlife Service Digital Library. Used by permission.

To aid conservationists in assessment of stream quality and evaluate the reliability of MIS and sentinel species at a local scale, with implications for larger regional and national areas, we investigated (1) the distribution of the Mottled Sculpin and the Slimy Sculpin in the upper Manistee River and its tributaries in the Manistee National Forest portion of the HMNF, (2) distributional and molecular differences between the two species, and (3) environmental features associated with the abundance of each species. The fundamental questions we addressed were: (1) can Mottled Sculpin be reliably distinguished from Slimy Sculpin?; (2) are distributions of these species in the HMNF sufficiently ubiquitous and sufficiently distinct for habitat assessment?; and (3) is the presence and abundance of Mottled Sculpin and Slimy Sculpin associated with stable abiotic stream conditions or with more complex metrics of stream quality scores (SQS)? Our results provide insights into the initial resolution of these questions.

## Materials and Methods

### Study sites and sampling procedures

We selected 12 sites in the Manistee River watershed (hereafter, primary sampling sites), beginning near the headwaters of the north branch and the main branch of the Manistee River in Kalkaska and Otsego counties, respectively, and progressing downstream through Crawford, Grand Traverse and Wexford counties (Fig 2, approximately 44° 29’ to 44° 54’ N, 84° 50’ to 85° 37’ W). Three sites were located on the north branch of the upper Manistee (M-72, Mecum Rd, and North Sharon Rd), six on the main branch of the upper Manistee (C-38, CD-1, CD-2, CD-3, Deward, and County Road 612), and three in tributary streams flowing into the upper Manistee (Little Cannon Creek, Manton Creek, and Anderson Creek). Sampled sites were 5–20 km apart within each branch or among tributaries. Each sampling reach was 50 m of main channel, a length we found sufficient to contain ≥2 geomorphic channel units [46, 47] or ≥2 different channel habitat units [48]. In addition to these 12 primary sampling sites along the Manistee River, we also obtained sculpin samples for molecular analyses from the nearby Au Sable, Boardman, Jordan, and Rapid Rivers, as well as from the Manistee River at its intersection with Cameron Bridge Road and near its mouth (Lower Manistee River) (hereafter, secondary sampling sites).

**Fig 2.**
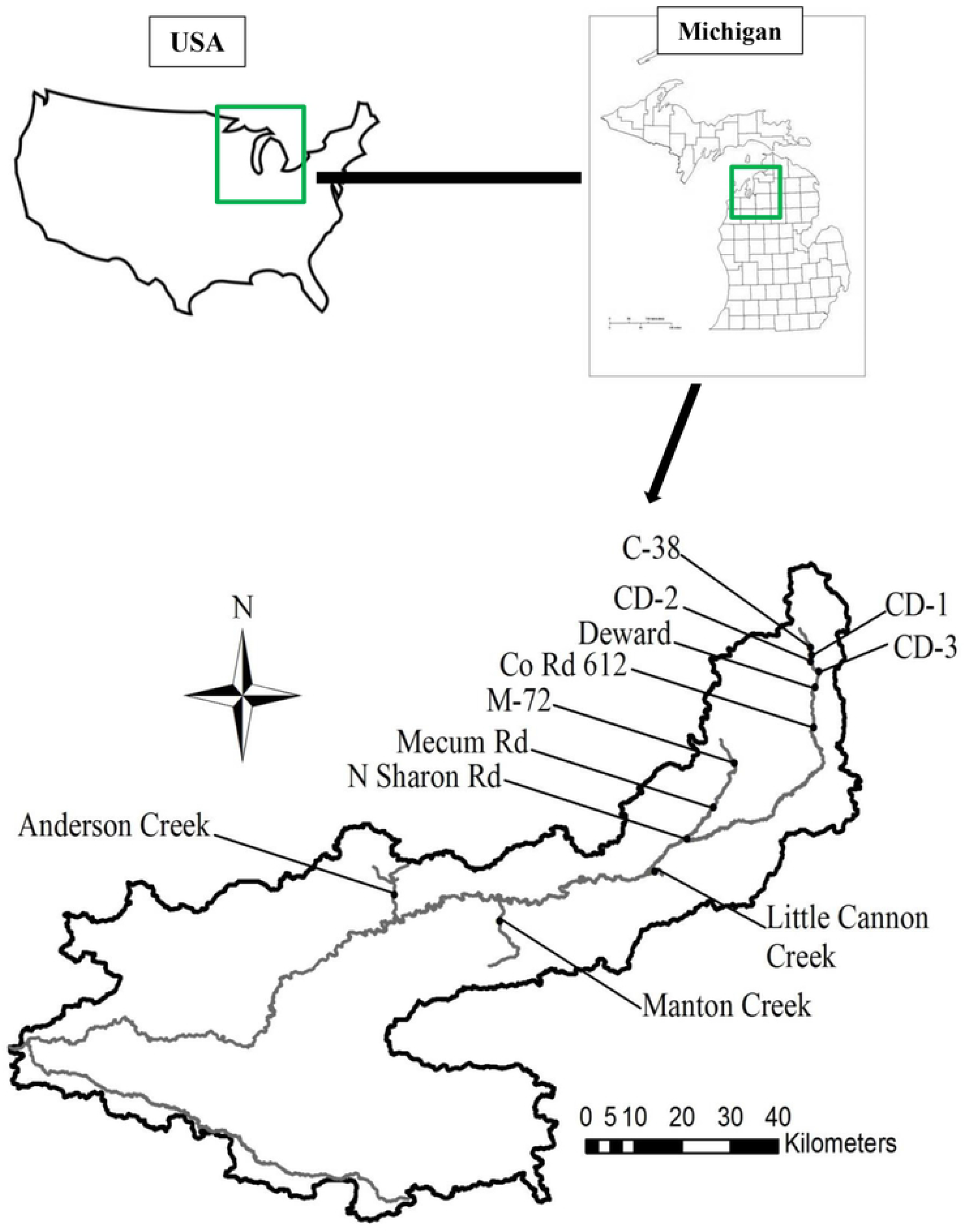
**Locations of 12 sampling sites in the Manistee River and its tributaries in Otsego, Crawford, Kalkaska, Wexford, and Grand Traverse Counties, Michigan (USA), approximately 44° 29’ to 44° 54’ N, 84° 50’ to 85° 37’ W**.

We collected sculpin with single-pass electrofishing during June and July 2015 using a Smith Root LR-24 Backpack Electrofisher with a voltage setting ranging between 200 to 280 V and a duty cycle set at a constant of 17 percent. We chose these months as periods providing the most stable summer conditions for Mottled and Slimy Sculpin and other aquatic species and less susceptible to temperature and discharge fluctuations common in both earlier and later periods. Collection time was recorded in seconds by the electrofisher during operation and then converted to minutes to compute sculpin abundance (sculpin/unit effort (min) = Catch Per Unit Effort (CPUE)) as an index of relative abundance. We collected and identified sculpin at each site with visual examination using a 10× hand lens. We selected ten sculpin individuals from each site and preserved them in ethanol (95%) for further assessment in the laboratory, the minimum number we considered necessary for more detailed morphological and DNA analysis and identification. We also collected northern redbelly dace (*Phoxinus eos* (Cope 1861)), Central mudminnow (*Umbra limi* (Kirtland 1841)), Brook trout (*Salvelinus fontinalis* (Mitchill 1814)), Brook stickleback (*Culaea inconstans* (Kirtland 1840)), and Round goby (*Neogobius melanostomus* (Pallas 1814)) in similar minimum numbers to serve as controls for DNA identification. The methods used in this study, specifically the capture and preservation of fish and macroinvertebrates, followed guidelines of the American Fisheries Society [49] and were reviewed and approved by the LRBOI Inland Fisheries Program, Manistee, Michigan USA. We followed these approved procedures in the field and laboratory.

### Morphological and molecular procedures

We used binocular dissecting microscopes to identify fish specimens in the laboratory using the key of Bailey et al. [44] (2004), which relies on the number of pelvic fin rays as the primary means to distinguish Mottled Sculpin from Slimy Sculpin. We then vouchered each sculpin into individual sample jars filled with ethanol (95%) for preservation. We clipped tail fins for DNA extraction conducted with the DNeasy Blood and Tissue Kit (Qiagen).

We amplified the Cytochrome Oxidase Subunit 1 (CO1) portion of the mitochondrial genome using the polymerase chain reaction (PCR). We used the primers VF2_1, FishF2_t1, FishR2_t1, and FR1d_t1 for amplification [50]. The expected bands for this primer combination are approximately 780 base pairs. We conducted PCR in 50 µl reactions consisting of 5 µl of 10× supplied buffer, 4 µl dNTP mixture (2.5 mM each), 1 µl Titanium *Taq* DNA Polymerase, 1 µl unquantified template DNA, 1 µl of each of the four primers (10 mM each), 35 µl of sterilized, distilled water, and a drop of mineral oil in a Perkin Elmer Cetus thermal cycler with 35 cycles of 94° C for 30 seconds, 52° C for 40 seconds, and 68° C for 60 seconds. To determine if amplification was successful, we loaded 5 µl of PCR product onto a precast FlashGel and electrophoresed it according to the manufacturer’s recommendations. We used ExoSAP-IT to enzymatically clean the amplified DNA. Functional Biosciences (Madison, WI) performed the capillary sequencing using primers M13F and M13R, which are embedded within the amplification primers [50].

We edited the chromatograms of the sequenced DNA using Sequencher v. 5.2.4 (GeneCodes, Ann Arbor, MI) and conducted a BLAST search of the GenBank database with each obtained sequence and used mmax score to identify species matches. We then aligned the sequences into a matrix using Mesquite v. 2.75 [51]. We analyzed this matrix phylogenetically using Maximum Parsimony and Distance methods in PAUP* v. 4.0a147 [52]. We used both the alignment visualized in Mesquite and the results of the preliminary phylogenetic analyses to identify samples with identical sequences. We verified differences between haplotypes, most of which consist of a single base pair difference, by consulting the chromatograms in Sequencher. Once the chromatograms had been reexamined to confirm each of the identified haplotypes, we excluded all but one sample of each haplotype in order to generate a matrix of the haplotypes and non-sculpin outgroups. We removed outgroups and constructed a Minimum Spanning Network [53] in PopART (Population Analysis with Reticulate Trees; http://popart.otago.ac.nz). We also analyzed the dataset in both a Distance and Maximum Parsimony framework in PAUP*. We conducted a heuristic search with Distance set as the criterion, and a full heuristic bootstrap search of 10,000 replicates, including groups compatible with the 50% majority-rule consensus, starting with stepwise addition, simple taxon addition, and TBR branchswapping.

### Sculpin and stream quality analyses

We recorded temperature (° C), dissolved oxygen (DO, mg/L), pH, conductivity (ms/cm), and turbidity (NTU) at all primary sampling sites with a Hydrolab HL4 (Hydrolab) (Supporting Document 1). We did not control for temperature variation relative to time of day or distance from headwaters, as the Manistee River is a groundwater-fed stream in which groundwater seepage influences temperature by providing a relatively cool input in summer and warm input in winter. In groundwater-fed streams, groundwater seepage affects provision of suitable water temperatures for aquatic biota and moderates other effects that could cause more extreme temperature changes [54]. In such streams, air temperature-stream temperature relationships are strongly related to and largely controlled by groundwater input [55]. We generated an estimate of elevation of each site by inserting GPS coordinates into Google Earth, and subsequently used elevation as an index of the site’s downstream distance in the watershed.

We collected macroinvertebrates at all primary sampling sites excluding M2 (CD-1), M3 (CD-2), and M4 (CD-3) (Fig 2). The sampling protocol followed procedures specified by Michigan Clean Water Corps [56]. We sorted macroinvertebrates to order and used their presence, abundance and diversity to compute a Stream Quality Score (SQS), a metric that uses presence and abundance of different orders of stream macroinvertebrates, based on their association with water quality and tolerance to pollution, to generate a numerical index that can be associated with various categories of stream conditions.

### Statistical analyses

We used Poisson regression to examine the relationship between elevation and CPUE for each sculpin species. For data modeling, values for site elevation were correspondingly shifted so that, to emphasize relative changes over the study area, designated elevation for the first site upstream was 0 m; in this regard, elevation difference between sites did not change in the analysis. For variables defining in-stream conditions, we generated boxplots of raw data relating Mottled Sculpin and Slimy Sculpin presence to temperature, DO, conductivity, pH, turbidity, and SQS. We conducted all analyses in R v. 4.0.5 [57].

## Results

### Genetic analysis, DNA sequencing, and haplotypes

Genetic analysis showed 87.5% correct laboratory identification of sculpin specimens relative to field (morphological) identification of sculpin collected during the first two weeks of the study, with errors confined to juvenile (<15 mm) specimens in which morphology was not yet fully developed. Field identifications based on morphology from the remaining field work were a 100% match with molecular determinations. BLAST searches successfully differentiated between Mottled Sculpin and Slimy Sculpin, supporting morphological distinction based on numbers of pelvic fin rays.

All fish samples for which DNA sequencing was attempted were successful. The CO1 sequences were edited to an aligned length of 671 basepairs, including two single basepair insertions in the sequence of one outgroup (*Umbra limi*). The alignment was otherwise completely unambiguous. Excluding outgroups, of the 669 basepairs obtained from the sculpin samples, 22 (3.3%) were variable and 18 (2.7%) were parsimony-informative.

All of the sculpin samples from which DNA was extracted yielded CO1 sequences that were adequate for assigning the sample to one or the other species. Of 121 DNA-barcoded sculpin samples, 60 were determined as Slimy Sculpin, and 61 as Mottled Sculpin. Twelve distinct CO1 haplotypes were recovered, five of Mottled Sculpin (H1–H5) and seven of Slimy Sculpin (H6–H12; Table 1). The CO1 sequence of each sculpin haplotype was deposited in GenBank, as well as those for the newly-generated outgroup CO1 sequences (Table 1). Of the 15 sites from which sculpins were sampled, five had only Mottled Sculpin, three only Slimy Sculpin, and seven had both species (Table 2). Seven sites contained a single CO1 haplotype Eight sites had between two and five haplotypes (Table 2).

**Table 1.** Sculpin haplotypes recovered, including the number of individuals of each haplotype and the sites at which they were found. The numbers in square brackets in the Haplotype column are the GenBank accession numbers.

| Haplotype | Species | Number of individuals recovered | Sites recovered |
| --- | --- | --- | --- |
| H1 [MW280589] | *Cottus bairdii* | 2 | 1 (M-72) |
| H2 [MW280588] | *Cottus bairdii* | 50 | 9 (C-38, County Rd 612, M-72, Mecum Rd, North Sharon Rd CD-1, CD-2, CD-3, Boardman River) |
| H3 [MW280590] | *Cottus bairdii* | 1 | 1 (County Rd 612) |
| H4 [MW280592] | *Cottus bairdii* | 2 | 1 (Lower Manistee River) |
| H5 [MW280591] | *Cottus bairdii* | 6 | 2 (Au Sable River, Lower Manistee River) |
| H6 [MW280597] | *Cottus cognatus* | 1 | 1 (CD-2) |
| H7 [MW280598] | *Cottus cognatus* | 1 | 1 (CD-2) |
| H8 [MW280593] | *Cottus cognatus* | 1 | 1 (Deward) |
| H9 [MW280595] | *Cottus cognatus* | 17 | 5 (Deward,<br>Cameron Bridge,<br>Jordan River,<br>CD-1, CD-2,<br>CD-3) |
| H10<br>[MW280599] | *Cottus cognatus* | 1 | 1 (CD-1) |
| H11<br>[MW280594] | *Cottus cognatus* | 38 | 6 (Deward,<br>County Rd 612,<br>CD-2, CD-3,<br>Boardman River,<br>Rapid River) |
| H12<br>[MW280596] | *Cottus cognatus* | 1 | 1 (Lower<br>Manistee River) |
| [MW280587] | *Culea inconstans* | 1 | Rapid River |
| [MW280584] | *Negobius*<br>*melanotomus* | 1 | Lower Manistee<br>River |
| [MW280586] | *Salvelinus*<br>*fontinalis* | 2 | Rapid River |
| [MW280585] | *Umbra limi* | 1 | Louie's Pond |

**Table 2.** Sculpin species and haplotypes represented at the collection sites. Sites C-38 through County Rd 612 were on the north branch of the Manistee River and are arranged in the table from upstream to downstream. Sites M-72 through North Sharon Road were on the main branch of the Manistee River, and are also arranged from upstream to downstream.

| Site | Species present | Haplotypes present |
| --- | --- | --- |
| C-38 | *Cottus bairdii* | 1 (H2) |
| CD-1 | *Cottus bairdii*, *C. cognatus* | 3 (H2, H9, H10) |
| CD-2 | *Cottus bairdii*, *C. cognatus* | 5 (H2, H6, H7, H9, H11) |
| CD-3 | *Cottus bairdii*, *C. cognatus* | 3 (H2, H9, H11) |
| Deward | *Cottus bairdii*, *C. cognatus* | 4 (H2, H8, H9, H11) |
| Cameron Bridge | *Cottus cognatus* | 1 (H9) |
| County Rd 612 | *Cottus bairdii*, *C. cognatus* | 3 (H2, H3, H11) |
| M-72 | *Cottus bairdii* | 2 (H1, H2) |
| Mecum Road | *Cottus bairdii* | 1 (H2) |
| North Sharon Road | *Cottus bairdii* | 1 (H2) |
| Au Sable River | *Cottus bairdii* | 1 (H5) |
| Boardman River | *Cottus bairdii*, *C.*<br>*cognatus* | 2 (H2, H11) |
| Rapid River | *Cottus cognatus* | 1 (H11) |
| Jordan River | *Cottus cognatus* | 1 (H9) |
| Lower Manistee<br>River | *Cottus bairdii*, *C.*<br>*cognatus* | 3 (H4, H5, H12) |

### Phylogenetic analyses

The Minimum Spanning Network we obtained from PopART (Fig 3) provided visual depiction of 11 basepairs that consistently differentiated Mottled and Slimy Sculpin. Each of the seven Slimy Sculpin haplotypes differed from their most similar haplotype by a single basepair. The maximum that any Slimy Sculpin haplotype differed from another was four basepairs (Fig 3). We found a similar pattern for Mottled Sculpin, except that two haplotypes, H2 and H5, differed by four basepairs (Fig 3), suggesting that, with further sampling, more Mottled Sculpin haplotypes might be recovered in the Manistee River. Both the Distance-based phylogeny and Maximum Parsimony-based bootstrap consensus tree (Fig 4) recovered Slimy and Mottled Sculpin haplotypes as reciprocally monophyletic groups, but while Slimy Sculpins were recovered with 99% bootstrap support, Mottled Sculpins were recovered with only 45% support (Fig 4).

**Fig 3.**
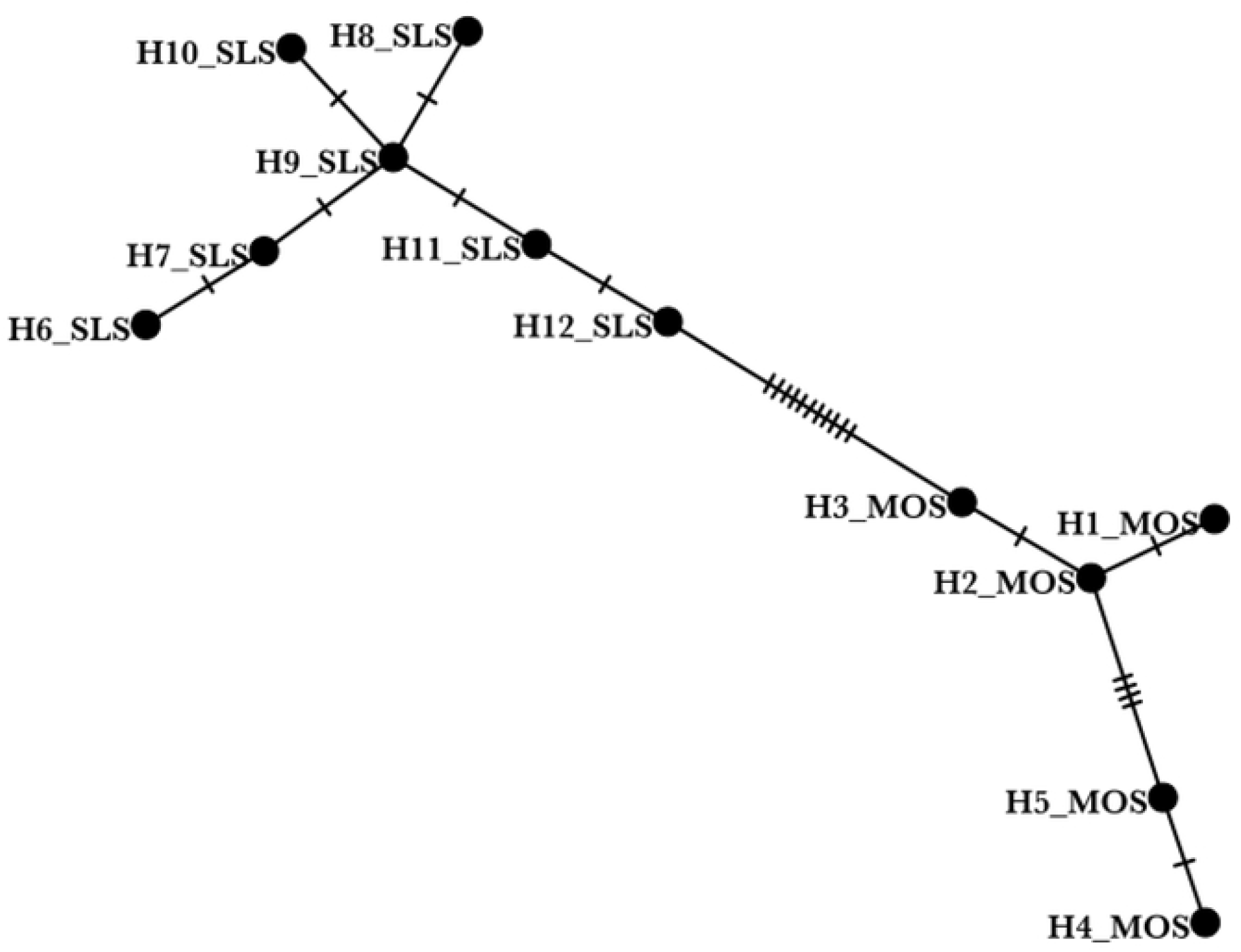
Minimum spanning network of the 12 Cottus haplotypes recovered from Northern Michigan. Each dash on the branches represents a base pair difference. Circles represent haplotypes, and are labeled with the haplotype number and “MOS” for Mottled Sculpin and “SLS” for Slimy Sculpin.

**Fig 4.**
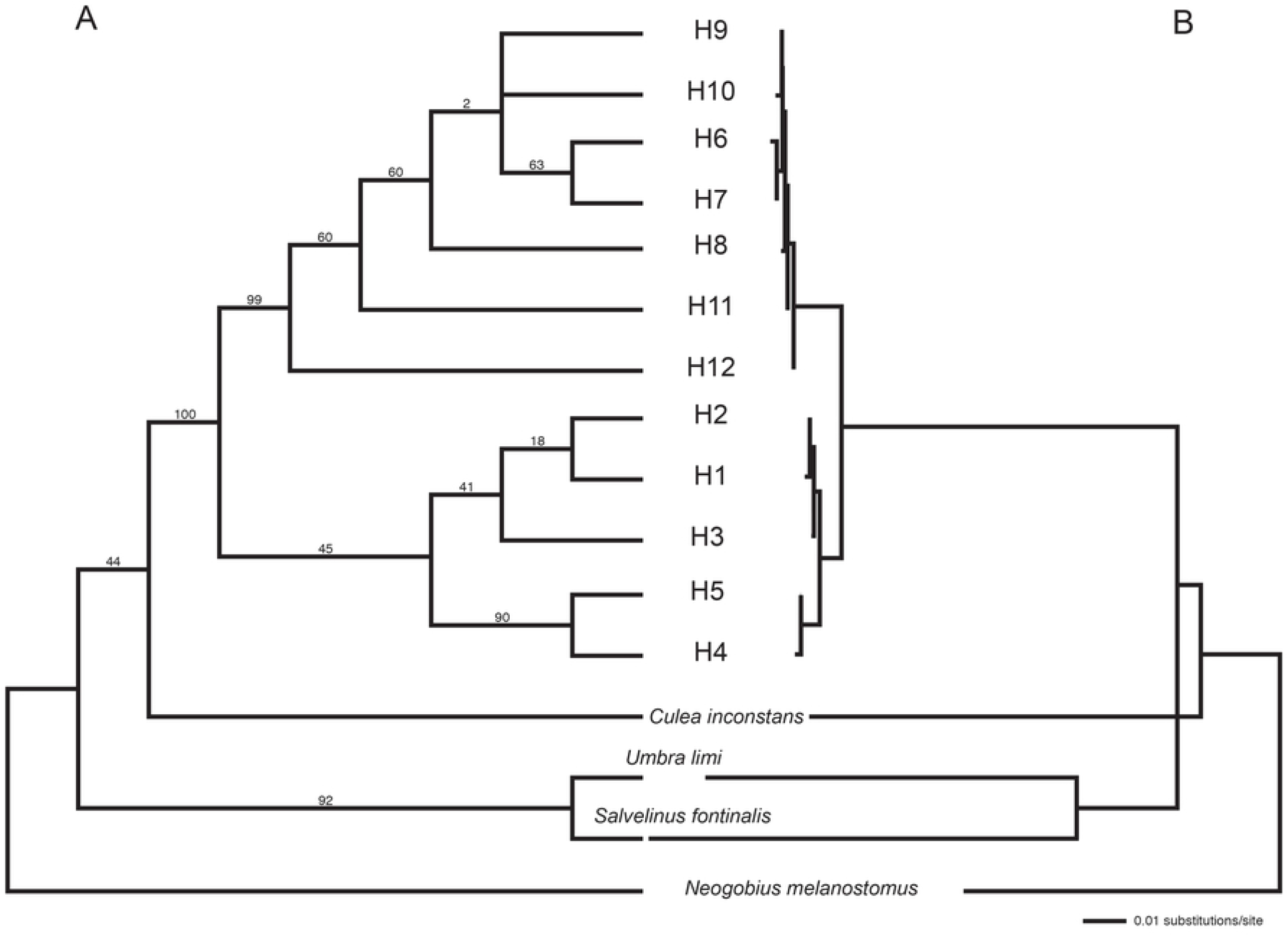
**A. Phylogram of 12 Cottus CO1 haplotypes and 4 outgroups of other fish species (Culea inconstans, Negobius melanotomus, Salvelinus fontinalis, and Umbra limi) computed in PAUP* with Distance as the criterion**. Branch lengths are proportional to the genetic distance along them. Numbers above branches are the substitutions per site estimated along each branch. Given the number of basepairs included in the analysis, a single basepair difference is equivalent to 0.001 substitutions per site on this phylogenetic tree. B. Maximum Parsimony bootstrap consensus tree of 10,000 bootstrap replicates of 12 Cottus CO1 haplotypes and 4 outgroups of other fish species (*Culea inconstans, Negobius melanotomus, Salvelinus fontinalis*, and *Umbra limi*) computed in PAUP*. Haplotypes are labeled with the haplotype number, haplotypes 1–5 are Mottled Sculpin, 6–12 are Slimy Sculpin.

### Factors affecting sculpin presence

Visual inspection of box plots showed that sites with Mottled Sculpin present had lower conductivity, pH, and stream quality and had considerably narrower range of colder temperatures than sites without Mottled Sculpin (Fig 5). In contrast, sites with Slimy Sculpin present had higher levels of DO and narrower ranges of conductivity, pH, temperature, and turbidity than sites without Slimy Sculpin (Fig 5).

**Fig 5.**
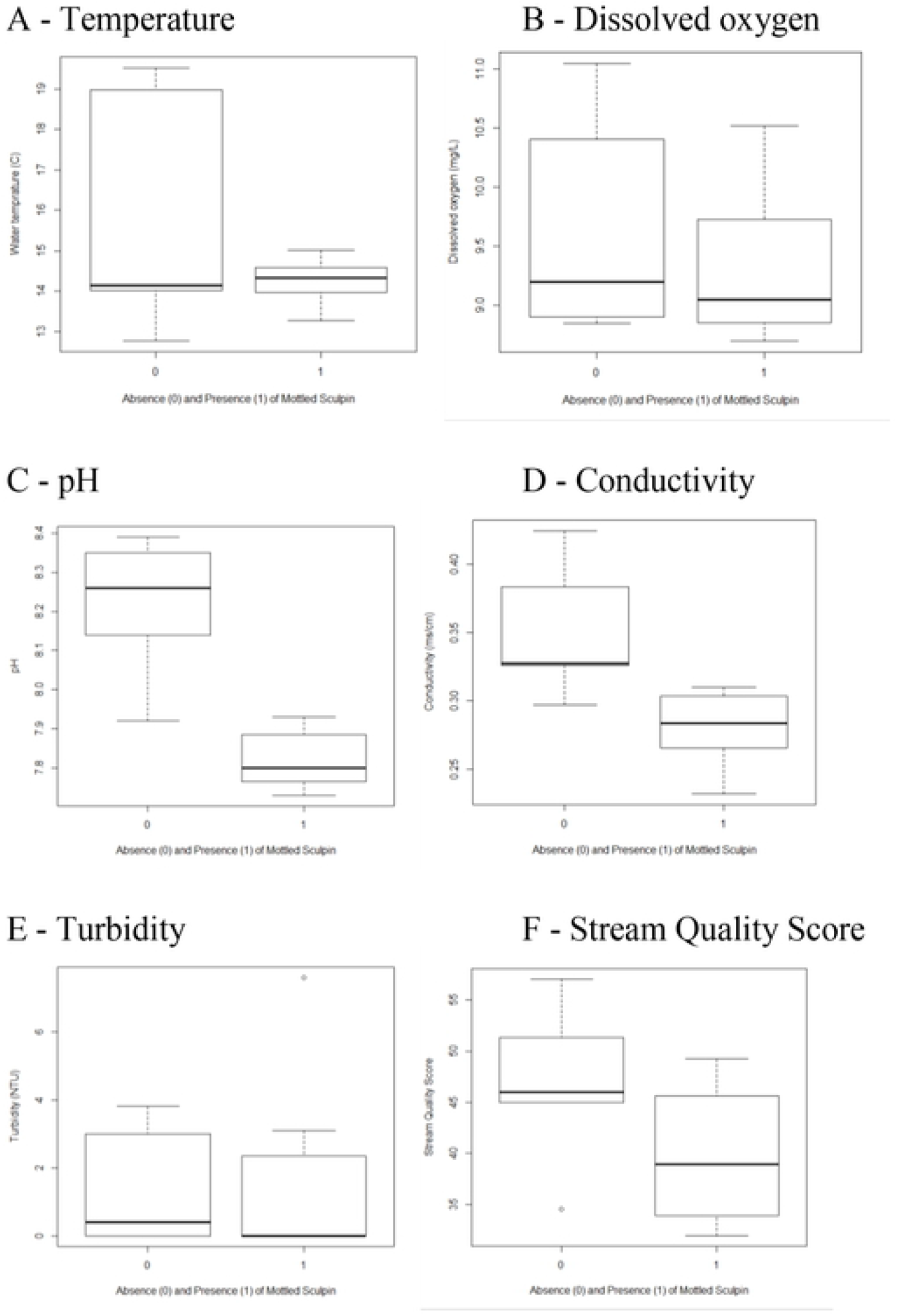
**Box plots of relationships between water quality parameters and absence or presence of Mottled Sculpin (Cottus bairdii) and Slimy Sculpin (C. cognatus) in the upper Manistee River, Michigan, USA**. Stream quality scores were based on orders of macroinvertebrates discovered, with the presence of less tolerant macroinvertebrates elevating the score. >48 = Excellent, 34-48 = Good, 19-33 = Fair, <19 = Poor (Michigan Clean Water Corps 2006). NTU refers to National Turbidity Units.

Abundance of both Mottled and Slimy Sculpin declined with elevation (i.e., increasing downstream distance), at the rate of 0.88 (0.74 – 0.96) Mottled Sculpin/min with each 1-m decline in elevation (*P* = 0.056) and for Slimy Sculpin 0.96 (0.86 – 1.15) less Slimy Sculpin/min with each 1-m decline in elevation (Fig 6). Over 10 km of stream length in the Manistee River, species-specific relative abundance, based on molecular identification, shifted from 100% Mottled Sculpin at C-38 and CD-1, the two most upstream sites, to a majority of Slimy Sculpin at the medium elevation sites, and again 100% Mottled Sculpin at the lowest elevation occupied sites (M-72, Mecum Rd, and N Sharon Rd). There were no sculpin of either species captured at the three lowest elevation sites (Supporting Document 1).

**Fig 6.**
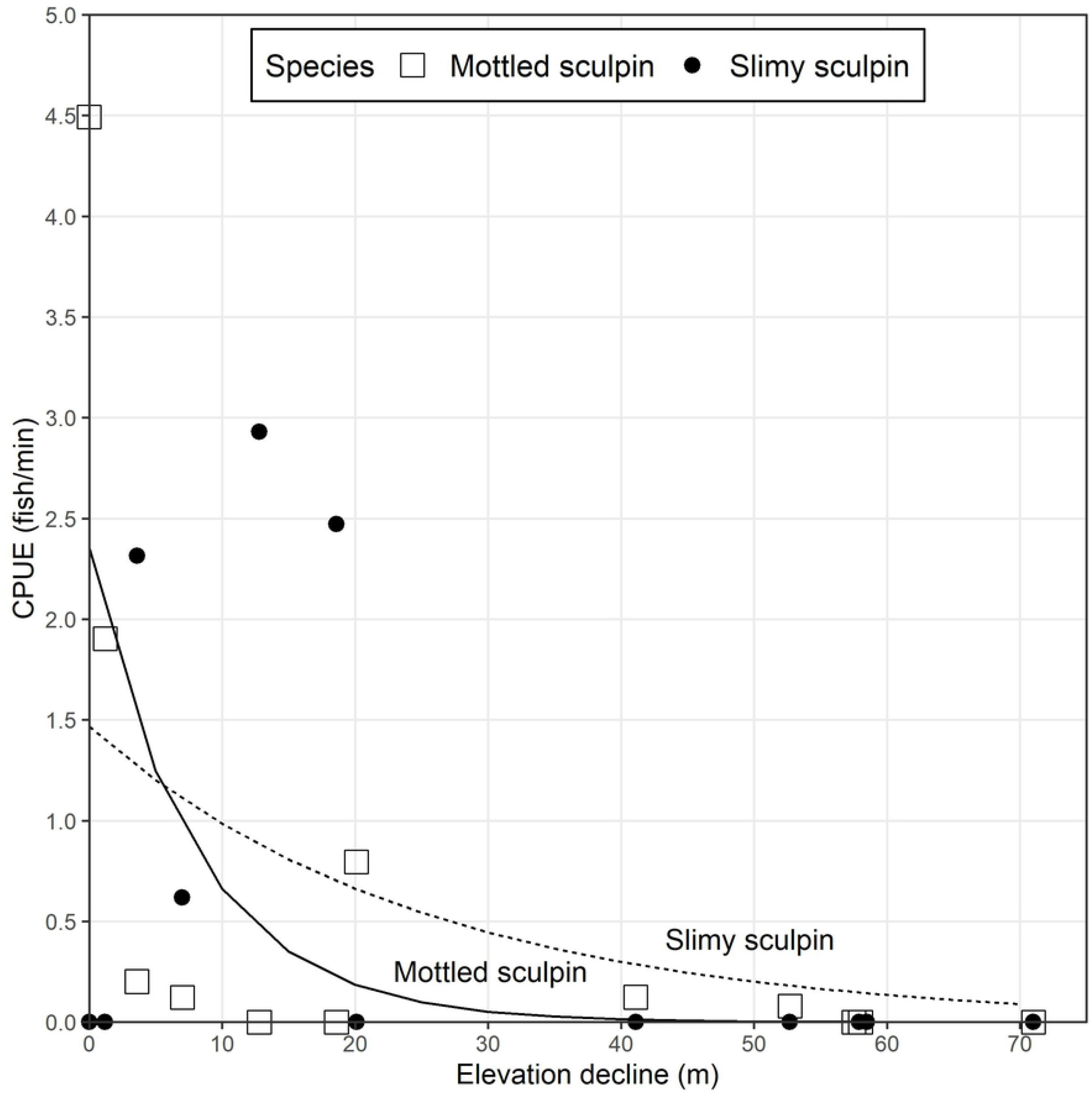
**Relationship (Poisson regression) between catch per unit effort (CPUE) and elevation decline (m) from highest elevation site for sites with Mottled Sculpin (Cottus bairdii) or Slimy Sculpin (C. cognatus) present in the Manistee River and its tributaries in Otsego, Crawford, Kalkaska, Wexford, and Grand Traverse Counties, Michigan, (USA)**

## Discussion

### Identification of Mottled Sculpin and Slimy Sculpin

Our first fundamental question was: can mottled sculpin be reliably distinguished from slimy sculpin? This is an important inquiry because species identifications are difficult in this genus. *Cottus* has long been recognized as taxonomically challenging, especially given the many proposals for new species classifications [58, 59] and ongoing debates as to phylogenetic relationships [60]. Problems in identification can be exacerbated by the frequent sympatry of sculpin species [61, 62]. Mottled Sculpin are sympatric with Potomac Sculpin (*C. girardi*) in Virginia (USA) [63, 64], Paiute Sculpin (*C. beldingi*) in Idaho and Wyoming (USA) [25], and Shoshone Sculpin (*C. greenei*) in Idaho [23]. Both Mottled and Slimy Sculpin are sympatric with Shorthead Sculpin (*C. confuses*) in western Canada [65, 66] and with one another in the western United States [25]. Challenges to identification are further compounded by the morphological similarity of different species, the tendency of different species to hybridize with one another [59, 67], and the proclivity to form new species in individual river systems [68]. Overlapping distribution contributed little to problems in species differentiation in our study, however, because the region of sympatry in the upper Manistee River was small. Nevertheless, identification based on physical characteristics alone remains suspect because of high morphological similarity.

Although problems in identification should not be minimized, we found that Mottled Sculpin could be distinguished from Slimy Sculpin using both morphological and molecular evidence, although, at the morphological level, sympatry of Mottled and Slimy Sculpin could be problematic to the value of using either species alone as an MIS or sentinel species. Where morphologically similar species are sympatric, risk of error in identification increases risk of error in conservation decisions. We found this risk to be low in the upper Manistee River and its principal tributaries, however, given that our genetic analyses confirmed a high degree of accuracy in correctly identifying adult sculpin in the field even with low-power microscopic examination, especially as we gained experience in identifying these species. Although we found differentiation between these two similar species to be relatively easy and reliable, it was best to observe individuals under a dissecting microscope, which we accomplished by preserving them in alcohol. Field determination, which would allow for release of the identified individuals, was slightly less reliable. Furthermore, the observed sympatry between Mottled Sculpin and Slimy Sculpin, although occurring only in a relatively small portion of the Manistee River’s total stream length, still offered opportunity for hybridization. Such hybridization could lead to ambiguity in identification, although we did not identify any individuals of obvious intermediate morphology, which would be one expectation if hybridization were occurring. This is relevant in the HMNF context because Slimy Sculpin have been shown to hybridize with Mottled Sculpin [68] and Rocky Mountain Sculpin, a species once classified in the *Cottus bairdii* complex [59]. Additional phylogenetic studies beyond the CO1 sequences generated for this study are needed to determine if hybrid sculpin are present in areas of sympatry [64].

Our own phylogenetic results obtained from the Minimum Spanning Network Distance and Maximum Parsimony suggested a down-to-up river phylogeographic pattern for the Slimy Sculpin, with haplotypes from further downstream being successively sister to more upstream haplotypes. The phylogeographic pattern for Mottled Sculpin was less obvious, but there was also indication that some Mottled Sculpin haplotypes were missing.

### Distribution of Mottled Sculpin and Slimy Sculpin

Our second fundamental question was: are distributions of the Mottled Sculpin and Slimy Sculpin in the HMNF sufficiently ubiquitous and sufficiently distinct for habitat assessment? We found that abundance of Mottled Sculpin and Slimy Sculpin declined with downstream distance throughout the Manistee River watershed, supporting the perception of both as headwater species. Such decline could be a factor of channel width, overall stream size, or other changes in hydrology associated with elevational change [59], but our data cannot address these hypotheses.

Distribution of Mottled Sculpin and Slimy Sculpin in the Manistee River was sympatric with respect to elevation, but allopatric with respect to individual sites. Regions of elevational sympatric distribution of Mottled Sculpin and Slimy Sculpin in portions of the Manistee suggest that some mechanism of niche separation may exist between the species. One mechanism that could exclude the Slimy Sculpin is the territorial behavior of Mottled Sculpin, which are known to compete against other species for feeding sites [69]. Additionally, geographic variation, water quality, and nonnative species might also contribute to niche separation in areas of overlap [25].

### Effects of environment on presence, absence, and abundance

Our third fundamental question was: was the presence and abundance of Mottled Sculpin and Slimy Sculpin associated with stable abiotic stream conditions or more complex metrics of stream quality scores? The relationships between environmental conditions and sculpin presence and abundance did not fully support the use of the Mottled Sculpin alone as an MIS for stream habitat quality, although the Mottled Sculpin was previously so designated in the HMNF. Both species preferred cold, clear waters, but Mottled Sculpin occurred in waters with lower conductivity, dissolved oxygen, and pH compared to Slimy Sculpin. The traditional conception of high-quality freshwater streams as that of streams with clear, cold, highly oxygenated water with expectedly diverse stream macroinvertebrate communities (and correspondingly high SQS) [70] matched some, but not all conditions associated with the presence of both species. Water temperature at sites with Mottled Sculpin was always cold, but sites where Mottled Sculpin were absent included both colder and warmer waters. Sites where Slimy Sculpin were present had higher levels of DO and lower levels of turbidity, and more constrained levels of conductivity, pH, temperature and turbidity than sampled sites in general.

Together the two species could provide sensitive indicators of all of these variables and thus function as a “sister species” version of an MIS. Imperfect associations between individual indicator species and the habitats or conditions they have been presumed to indicate can be routinely expected [71]. Similarly, other investigators have shown that sculpin are sensitive to many variables indicative of water conditions to a varying, and not always predictable, extent [25]. We found Mottled Sculpin only in streams with turbidity, temperature, and DO at levels characteristic of high-quality streams, but did not find Mottled Sculpin in every high quality stream we sampled (Supporting Document 1). Slimy Sculpin were also found in cold water streams with similar stream quality scores, but otherwise displayed sensitivity to different variables. This distributional arrangement suggests that these species might be indicators of higher quality stream conditions. However, if headwater habitats are periodically colonized by both Mottled Sculpin and Slimy Sculpin, then patch dynamics might produce variability in abundance without meaningful differences in habitat quality.

Species associations with particular sites or habitats are not necessarily indicators that the site or habitat reflects the niche of the species. As De Cáceres et al. [17] point out, such associations may be (1) a random event, (2) reflections of historical events (environmental perturbations in the system) or non-niche-related species characteristics (population fluctuations or dispersal patterns), or (3) genuine associations reflecting the species preferences, but not ones the manager is assuming or even aware of. The third problem may be reflected in factors we did not examine in this study, including and especially habitat structure, which might be more important than water conditions. For example, Bond and Jones [72] noted the importance of longitudinal gradients in rivers with respect to fish abundance and distribution, with longitude serving as an integrator of changes in velocity, temperature, food resources (including and especially macroinvertebrates), and coarse particulate matter. Quist et al. [25] determined that Mottled Sculpin were absent from reaches in mountain streams with high velocity and large rocky substrates, suggesting that they required slower-water habitats for a portion of their development. Similarly, some streams in the Manistee River watershed that did not contain both species might be healthy streams in which one species was absent because the stream did not provide sufficient habitat juxtaposition for adequate niche separation. Such findings might explain why no sculpin were found at our three most downstream sites (Little Cannon Creek, Manton Creek, and Anderson Creek), despite those sites possessing similar water quality in most variables and higher stream quality scores than other sites.

We cannot offer an explanation of why Mottled Sculpin were present at sites with lower SQS scores, which are primarily reflections of composition and diversity of the stream macroinvertebrate community, a metric that many view as the single best biotic indicator of stream condition [4]. Other studies have noted that the relationship between sculpin and macroinvertebrates is not a correlation between independent assessments of water quality, but an expression of trophic linkage [73]. Macroinvertebrate prey abundance influences patch selection by sculpin more than physical microhabitat characteristics [74], but dynamics of macroinvertebrate populations and habitat structure influencing sculpin distribution are not well understood, nor were they examined in our study. It is possible that sculpin abundance in the Manistee River and similar streams does not correlate with macroinvertebrates in ways related to SQS, but to specific distributions and abundances of individual macroinvertebrate species.

## Conclusions

Our data were limited to a single river, and therefore cannot be assumed to apply to other watersheds without careful study and qualification. Nevertheless, we offer some general insights from this investigation that might have wider applicability. Rather than using only Mottled Sculpin as an MIS, as it has been designated, we recommend an MIS complex of Mottled Sculpin and Slimy Sculpin. Using Mottled Sculpin alone could lead to disregard of high-quality waters where this species is not present, and Mottled Sculpin lack a sufficiently ubiquitous distribution to ignore this risk. Inclusion of the Slimy Sculpin would permit coverage of a greater proportion of the watershed, as well as reduce concerns regarding incorrect identification at the species level given that such distinction would not need to be made. Additionally, and alternatively, there is warrant to consider using SQSs, such as those generated by estimation of stream macroinvertebrate diversity, instead of employing an MIS approach, and to incorporate consideration of variables affecting stream habitat structure, as well as water conditions. At a more systemic scale, managers should continue to refine and increasingly use quantitative models of species-site associations to, whenever possible, develop more precise understanding of underlying causative factors of such associations and what such associations actually indicate.

## Acknowledgments

Funding was provided by the Little River Band of Ottawa Indians, a Native American nation indigenous to Michigan (USA), as part of their efforts in research and rehabilitation of the Lake Sturgeon (*Accipiter fulvescens*), a species of high cultural significance to the Tribe. The findings, opinions, and recommendations expressed within this publication are those of the authors and not necessarily those of the Bureau of Indian Affairs (BIA) or of the Tribe. C. Riley, USFS, provided suggestions, ideas, and guidance in the development of the original research proposal. M. Holtgren, Inland Fisheries Program, Little River Band of Ottawa Indians, provided support, assistance, training in field techniques and encouragement throughout the study. Staff of the Au Sable Institute (Michigan, USA) provided room, board, and support for the research team. H. W. Garris, M. Holtgren, and R. Westerhof reviewed preliminary drafts of the manuscript.

## References

1. Dos Santos DA, Molineri C, Reynaga MC, Basualdo C. Which index is best to assess stream health? Ecological Indicators. 2011; 11:582–589.

2. Karr JR. Assessment of biotic integrity using fish communities. Fisheries. 1981; 6:21–27. doi.org/10.1577/1548-8446(1981)006<0021:AOBIUF>2.0.CO;2

3. Pander J, Geist J. Ecological indicators for stream restoration success. Ecological Indicators. 2013; 30:106–118. doi.org/10.1016/j.ecolind.2013.01.039

4. Lydy, MJ, Crawford CG, Frey JW. A comparison of selected diversity, similarity, and biotic indices for detecting changes in benthic-invertebrate community structure and stream quality. Archives of Environmental Contamination and Toxicology. 2000; 39:469–479. doi.org/10.1007/s002440010129

5. Mostafavi H, Schinegger R, Melcher A, Moder K, Mielach C, Schmutz S. A new fish-based multi-metric assessment index for cyprinid streams in the Iranian Caspian Sea. Basin. Limnologica. 2015; 51:37–52. doi.org/10.1016/j.limno.2014.10.006

6. Dufrêne M, Legendre P. Species assemblages and indicator species: The need for a flexible asymmetrical approach. Ecological Monographs. 1997; 67:345–366. doi.org/10.1890/0012-9615(1997)067[0345:SAAIST]2.0.CO;2

7. Bakker, JD. Increasing the utility of Indicator Species Analysis. J Applied Ecology. 2008; 45:1829–1835. doi: 10.1111/j.1365-2664.2008.01571.x

8. Cuffney, TF, Qian SS. A critique of the use of indicator-species scores for identifying thresholds in species responses. Freshwater Sci. 2013; 32:471-488. doi:10.1899/12-056.1

9. Dziock F, Henle K, Foeckler F, Follner K, Scholz M. Biological indicator systems in floodplains – a review. International Rev Hydrobiology. 2006; 4:271–291. doi.org/10.1002/iroh.200510885

10. Neumann M, Liess M, Schulz R. An expert system to estimate the pesticide contamination of small streams using benthic macroinvertebrates as bioindicators, Part 2:The knowledge base of LIMPACT. Ecological Indicators. 2003; 2:391–401. doi.org/10.1016/S1470-160X(03)00025-6

11. Bellinger EG, Sigee DC. Freshwater Algae: Identification and Use as Bioindicators. West Sussex: John Wiley & Sons; 2010.

12. Salovaara KJ, Cardenas GG, Tuomisto H. Forest classification in an Amazonian rainforest landscape using pteridophytes as indicator species. Ecography. 2004; 27:689–700. doi.org/10.1111/j.0906-7590.2004.03958.x

13. Lindenmayer DB, Margules CR, Dotkin DB. Indicators of biodiversity for ecologically sustainable forest management. Conservation Biol. 2000; 14:941–950. doi.org/10.1046/j.1523-1739.2000.98533.x

14. Van Dyke F. Conservation biology: foundations, concepts, applications. 2nd ed. Dordrecht: Springer; 2008.

15. Zacharias, MA, Roff XC. Use of focal species in marine conservation and management: a review and critique. Marine and Freshwater Ecosystems. 2001; 11:59–76.

16. Landres, PB, Verner J, Thomas JW. Ecological uses of vertebrate indicator species: a critique. Conservation Biol. 1988; 2:316–328. doi.org/10.1111/j.1523-1739.1988.tb00195.x

17. De Cáceres M, Legendre R, Moretti M. Improving indicator species analysis by combining groups of sites. Oikos. 2010; 119:1674–1684. doi.org/10.1111/j.1600-0706.2010.18334.x

18. Gray MA, Curry RA, Arciszewski TJ, Munkittrick KR, Brasfield SM. The biology and ecology of the slimy sculpin: a recipe for effective environmental monitoring. FACETS. 2018; 3:5 February. doi.org/10.1139/facets-2017-0069)

19. Weinstein BG, Graham CH, Parra JL. The role of environment, dispersal and competition in explaining reduced co-occurrence among related species. PLOS ONE. 2017; doi.org/10.1371/journal.pone.0185493.

20. NatureServe Explorer 2.0. Cottus bairdii Mottled Sculpin. 2021; Available from: https://explorer.natureserve.org/Taxon/ELEMENT_GLOBAL.2.819868/Cottus_bairdii

21. United States Geological Survey. Cottus bairdii Mottled Sculpin. 2021; Available from: https://explorer.natureserve.org/Taxon/ELEMENT_GLOBAL.2.819868/Cottus_bairdii

22. Besser JM, Mebane CA, Mount DR, Ivey CD, Kunz JL, Greer IE, May TW, Ingersoll CG. Sensitivity of Mottled Sculpins (Cottus bairdii) and Rainbow Trout (Onchorhynchus mykiss) to acute and chronic toxicity of Cadmium, Copper, and Zinc. Environmental Toxicology and Chemistry. 2007; 26:1657–65. doi: 10.1897/2006-571.S1

23. Kuda DB, Griffith JS. Establishment of Shoshone Sculpin (Cottus greenei) in a spring inhabited by Mottled Sculpin (C. bairdi). Great Basin Naturalist. 1993; 53:190–193.

24. Markle DF, Hill DL Jr. Taxonomy and distribution of the Malheur mottled sculpin, Cottus bendirei. Northwest Sci. 2000; 74:202–211. Quist MC, Hubert WA, Isaak DJ. Factors affecting allopatric and sympatric occurrence of two sculpin species across a Rocky Mountain watershed. Copeia. 2004; 3:617–623. doi.org/10.1643/CE-04-026R1

25. Quist MC, Hubert WA, Isaak DJ. Factors affecting allopatric and sympatric occurrence of two sculpin species across a Rocky Mountain watershed. Copeia. 2004; 3:617–623. https://doi.org/10.1643/CE-04-026R1

26. Petty JT, Grossman GD. Restricted movement by Mottled Sculpin (pisces:cottidae) in a southern Appalachian stream. Freshwater Biol. 2004; 49:631–645. doi.org/10.1111/j.1365-2427.2004.01216.x

27. Keeler RA, Breton AR, Peterson DP, Cunjak RA. Apparent survival and detection estimates for PIT-tagged slimy sculpin in five small New Brunswick streams. Trans American Fisheries Soc. 2007; 136(1):281–292. doi: 10.1577/T05-131.1

28. Breen MJ, Ruetz CR III, Thompson KJ, Kohler SL. Movements of Mottled Sculpins (Cottus bairdii) in a Michigan stream: how restricted are they? Canadian J Fisheries and Aquatic Sci. 2009; 66:31–41. doi: 10.1139/F08-189

29. Symons PEK, Metcalfe JL, Harding GD. Upper lethal and preferred temperatures of the slimy sculpin, Cottus cognatus. J. Fisheries Res Board of Canada. 1976; 33:180–183. doi: 10.1139/f76-022

30. Otto RG, Rice JO. Responses of a freshwater sculpin (Cottus cognatus gracilis) to temperature. Trans American Fisheries Soc. 1977; 106:89–94. doi: 10.1577/1548-8659(1977)106<89:ROAFSC>2.0.CO;2

31. Waite IR, Carpenter KD. Associations among fish assemblage structure and environmental variables in Willamette Basin streams, Oregon. 2000; Trans American Fisheries Soc. 129:754–770. doi.org/10.1577/1548-8659(2000)129<0754:AAFASA>2.3.CO;2

32. Lessard JL, Hayes DB. Effects of elevated water temperature on fish and macroinvertebrate communities below small dams. River Res Applications. 2003; 19:721–732. doi.org/10.1002/rra.713

33. Adams SB, Schmetterling DA. Freshwater sculpins: phylogenetics to ecology. Trans American Fisheries Soc. 2007:136:1736–1741. doi: 10.1577m7-023.1

34. Dubé MG, MacLatchy DL, Kieffer JD, Glozier NE, Culp JM, Cash KJ. Effects of metal mining effluent on Atlantic salmon (Salmo salar) and slimy sculpin (Cottus cognatus): using artificial streams to assess existing effects and predict future consequences. Sci. Total Environment. 2005; 343:135–154. doi: 10.1016/j.scitotenv.2004.09.037

35. Allert AL, Fairchild JF, Schmitt CJ, Besser JM, Brumbaugh WG, Olson SJ. Effects of mining-derived metals on riffle-dwelling benthic fishes in Southeast Missouri, USA. Ecotoxicology and Environmental Safety. 2009; 72(6):1642–1651. PMID: 19570577 doi: 10.1016/j.ecoenv.2009.02.014

36. Gray MA, Curry RA, Munkittrick KR. Non-lethal sampling techniques for assessing environmental impacts using a small-bodied sentinel fish species. Water Quality Res J Canada. 2002; 37:195–211. doi.org/10.2166/wqrj.2002.012

37. Brasfield SM, Hewitt LM, Chow L, Batchelor S, Rees H, Xing Z Z, et al. Assessing the contribution of multiple stressors affecting small-bodied fish populations through a gradient of agricultural inputs in northwestern New Brunswick, Canada. Water Quality Res. J. Canada. 2015; 50:182–197. doi: 10.2166/wqrjc.2014.126

38. Miller LL, Isaacs MA, Martyniuk CJ, Munkittrick KR. Using molecular biomarkers and traditional morphometric measurements to assess the health of slimy sculpin (Cottus cognatus) from streams with elevated selenium in North-Eastern British Columbia. Environmental Toxicology and Chemistry. 2015; 34(10):2335–46. PMID: 25982233 doi: 10.1002/etc.3064

39. Galloway BJ, Munkittrick KR, Currie S, Gray M, Curry RA, Wood CS. Examination of the responses of slimy sculpin (Cottus cognatus) and white sucker (Catostomus commersoni) collected on the Saint John River (Canada) downstream of pulp mill, paper mill, and sewage discharges. Environmental Toxicology and Chemistry. 2003; 22:2898–2907. PMID: 14713029 doi: 10.1897/02-181

40. Arciszewski TJ, Kidd KA, Munkittrick KR. Comparing responses in the performance of sentinel populations of stoneflies (Plecoptera) and slimy sculpin (Cottus cognatus) exposed to enriching effluents. Ecotoxicology and Environmental Safety. 2011; 74:1844–1854. PMID: 21816476. doi: 10.1016/j.ecoenv.2011.07.010

41. U. S. Forest Service (USFS). Huron Manistee Forest Plan and Environmental Impact Statement. Appendix G. 2013. Available from: http://www.fs.usda.gov/Internet/FSE_DOCUMENTS/fsm8_046666.pdf

42. Resetarits WJ Jr. Limiting similarity and the intensity of competitive effects on the Mottled Sculpin, Cottus bairdii, in experimental stream communities. Oecologia. 1995; 104:31–38. doi.org/10.1007/BF00365559

43. McAllister DE. Distinguishing characteristics for the sculpins Cottus bairdi and Cottus cognatus in eastern Canada. J Fisheries Res Board Canada. 1964; 21:1339–1342. doi: 10.1139/f64-113

44. Bailey RM, Latta WC, Smith GR. An atlas of Michigan fishes with keys and illustrations for their identification. Miscellaneous Publications. Museum of Zoology. No. 192. Ann Arbor: University of Michigan; 2004.

45. Baker RL, Chandler LM, Eckdhal TT. Identification of Cottus species in Montana using mitochondrial RFLP analysis. Bios. 2001; 72:87–91.

46. Lyons J. The length of stream to sample with a towed electrofishing unit when fish species richness is estimated. North American J Fisheries Management. 1992; 12:198–203. doi.org/10.1577/1548-8675(1992)012<0198:TLOSTS>2.3.CO;2

47. Meador MR, Cuffney TF, Gurtz ME. Methods for sampling fish communities as part of the national water-quality assessment program. U. S. Geological Survey. U. S. Department of the Interior. Washington, D. C. 1993. Available from: https://water.usgs.gov/nawqa/protocols/OFR-93-104/fish5.html

48. Bisson PA, Montgomery DR, Buffington JM. Pages 21-47 in F. R. Hauer and G. Lambert, editors. Valley segments, stream reaches, and channel units. Methods in Stream Ecology 3rd edition. San Diego: Elsevier; 2017.

49. Use of Fishes in Research Committee (joint committee of the American Fisheries Society, the American Institute of Fishery Research Biologists, and the American Society of Ichthyologists and Herpetologists). Guidelines for the use of fishes in research. Bethesda: American Fisheries Soc.; 2014.

50. Ivanova NV, Zemlak TS, Hanner RH, Hebert PDN. Universal primer cocktails for fish DNA barcoding. Mol Ecology Notes. 2007; 7:544–548. doi.org/10.1111/j.1471-8286.2007.01748.x

51. Maddison D, Maddison W. Mesquite: a modular system for evolutionary analysis. 2011; Available from http://mesquiteproject.org

52. Swofford DL. PAUP*. Phylogenetic analysis using parsimony (* and other methods.) Sunderland: Sinauer Associates; 2002.

53. Bandelt H, Forster P, Röhl A. Median-joining networks for inferring intraspecific phylogenies. Mol Biol and Evolution. 1999:16:37–48. doi: 10.1093/oxfordjournals.molbev.a026036

54. Kaandorp VP, Doornenbal PJ, Kooi K, Broers HP, de Louw GB. Temperature buffering by groundwater in ecologically valuable lowland streams under current and future climate conditions. J Hydrology. 2019; 3. doi.org/10.1016/j.hydroa.2019.100031

55. Driscoll MO, Dewalle DR. Stream-air temperature relations to classify stream-groundwater interactions in a karst setting, central Pennsylvania, USA. J Hydrology. 2006; 329:140–153. doi.org/10.1016/j.jhydrol.2006.02.010

56. Michigan Clean Water Corps. MiCorps volunteer stream monitoring procedures. Michigan Clean Water Corps. 2006; Available from: https://micorps.net/wp-content/uploads/2021/01/VSMP-MonitoringProcedures.pdf

57. R Development Core Team. R: A language and environment for statistical computing. Version 4.0.5. Vienna: R Foundation for Statistical Computing; 2021.

58. Neely DA, Williams JD, Mayden RL. Two new sculpins of the genus Cottus (Teleostei: Cottidae) from rivers of North America. Copeia. 2007; 3:641–656. doi.org/10.1643/0045-8511(2007)2007[641:TNSOTG]2.0.CO;2

59. Rudolfsen TJ, Ruppert WL, Taylor EB, Davis CS, Watkinson DA, Poesch MS. Habitat use and hybridization between Rocky Mountain Sculpin (Cottus sp.) and Slimy Sculpin (Cottus cognatus). Freshwater Biol. 2019; 64:391–404. doi.org/10.1111/fwb.13225

60. Yokoyama R, Goto A. Evolutionary history of freshwater sculpins, genus Cottus (Teleostei: Cottidae) and related taxa, as inferred from mitochondrial DNA phylogeny. Mol Phylogenetics and Evolution. 2005. 36:654–668. doi.org/10.1016/j.ympev.2005.06.004

61. Mason JC, Machidori S. Populations of sympatric sculpins, cottus aleut/cus and cottus asper, in four adjacent salmon-producing coastal streams on Vancouver Island, B.C. Fishery Bull. 1976; 74:131–141.

62. Daniels, R. A. Comparative life histories and microhabitat use in three sympatric sculpins (Cottidae: Cottus) in northeastern California. Environmental Biol Fishes. 1987; 19:93–110. doi.org/10.1007/BF00001880

63. Matheson RE Jr, Brooks GR Jr. Habitat segregation between Cottus bairdi and Cottus Girardi: an example of complex inter- and intraspecific resource partitioning. American Midland Naturalist. 1983;110:165–176. doi.org/10.2307/2425222

64. Kinziger A, Raesly R. A narrow hybrid zone between two Cottus species in Wills Creek, Potomac Drainage. American Genet Assoc. 2001; 92:309–314. doi.org/10.1093/jhered/92.4.309

65. Hughes GW, Peden AE. Life history and status of the shorthead sculpin (Cottus confusus: Pisces, Cottidae) in Canada and the sympatric relationship to the slimy sculpin (Cottus cognatus). Canadian J Zoology. 1984; 62:306–311. doi.org/10.1139/z84-047

66. Peden AE, Hughes GW, Roberts WE. Morphologically distinct populations of the Shorthead Sculpin, Cottus confusus, and Mottled Sculpin, Cottus bairdi (Pisces, Cottidae), near the western border of Canada and the United States. Canadian J Zoology. 1989; 67:2711–2720. https://doi.org/10.1139/z89-384

67. Strauss RE. Natural hybrids of the freshwater sculpins Cottus bairdi and Cottus cognatus (Pisces: Cottidae): Electrophoretic and Morphometric Evidence. American Midland Naturalist. 1986; 115:87–105. doi.org/10.2307/2425839

68. Lemoine ML, Young MK, McKelvey KS, Eby L, Pilgrim KL, Schwartz MK. Cottus schitsuumsh, a new species of sculpin (Scorpaeniformes: Cottidae) in the Columbia River basin, Idaho-Montana, USA. 2014. doi:http://dx.doi.org/10.11646/zootaxa.3755.3.3

69. Petty JT, Grossman GD. Size-dependent territoriality of Mottled Sculpin in a southern Appalachian stream. Trans American Fish Soc. 2007; 136:1750–1761. doi.org/10.1577/T06-034.1

70. Environmental Protection Agency. Examples of watershed assessments for watershed health. USEPA. 2019. Available from: https://www.epa.gov/hwp/examples-water-quality-assessments-watershed-health

71. Barry D, Fischer RA, Hoffman KW, Barry T, Zimmerman EG, Dickson KL. Assessment of habitat values for indicator species and avian communities in a riparian forest. Southeastern Naturalist. 2006; 5:295–310. doi: 10.1656/1528-7092(2006)5[295:AOHVFI]2.0.CO;2

72. Bond MJ, Jones NE. Spatial distribution of fishes in hydropeaking tributaries of Lake Superior. River Research and Applications. 2015; 31:120–33. doi: 10.1002/rra.2720

73. McGinley E, Raesley R, Seddon W. The effects of embeddedness on the seasonal feeding of Mottled Sculpin. American Midland Naturalist. 2013; 170:213–228. doi.org/10.1674/0003-0031-170.2.213

74. Petty JT, Grossman GD. Patch selection by Mottled Sculpin (Pisces: Cottidae) in a southern Appalachian stream. Freshwater Biol. 1996; 35:261–76. https://doi.org/10.1046/j.1365-2427.1996.00498.x

